# Temperature-dependent ligand relocation reveals plasticity of TRPM4 inhibition

**DOI:** 10.64898/2026.05.13.724805

**Authors:** Dominic Schneiter, Jean-Sébastien Rougier, Hugues Abriel, Henning Stahlberg, Babatunde Ekundayo

**Affiliations:** Institute of Biochemistry and Molecular Medicine (IBMM), University of Bern, 3012 Bern, Switzerland; Laboratory of Biological Electron Microscopy (LBEM), Institute of Physics, School of Basic Sciences, Federal Institute of Technology Lausanne (EPFL), 1015 Lausanne, Switzerland; Department of Fundamental Microbiology, Faculty of Biology and Medicine, University of Lausanne, 1015 Lausanne, Switzerland

## Abstract

Transient receptor potential melastatin 4 (TRPM4) is a Ca²⁺-activated cation channel whose pharmacology is shaped by its molecular environment. It remains poorly understood how temperature and membrane context influence inhibitor recognition. Here we combine cryo-electron microscopy of membrane-derived vesicles and detergent-solubilized TRPM4 to investigate lipid-associated architecture and binding of the potent anthranilic anilide inhibitor PBA. We find that membrane vesicles preserve a native-like paralipid environment and reveal lipid binding patterns highly similar to those observed in GDN, supporting detergent-solubilized TRPM4 as a structurally relevant system for ligand analysis. Strikingly, PBA occupies distinct binding pockets at 8 °C and 37 °C. At low temperature, PBA binds in a previously described inhibitor pocket formed by S3, S4, the S4–S5 linker and the TRP helix, whereas at physiological temperature it relocates to a distinct site within the S1–S4 domain proximal to the Ca²⁺ regulatory region. These findings reveal temperature-dependent plasticity in TRPM4 ligand recognition.

## Introduction

Transient receptor potential melastatin 4 (TRPM4) is a Ca²⁺-activated, monovalent-selective cation channel that is broadly expressed in cardiac, neuronal, epithelial and immune tissues, where it couples elevations in intracellular Ca²⁺ to membrane depolarisation and thereby modulates diverse physiological processes^1,2^. TRPM4 dysfunction has been linked to inherited cardiac conduction disorders, Brugada syndrome and cancer-associated phenotypes, which has made the channel an increasingly attractive pharmacological target^2–8^.

Structural and functional studies have established TRPM4 as a highly dynamic channel regulated by intracellular Ca²⁺, membrane lipids and small molecules. Early cryo-electron microscopy (cryo-EM) studies defined the overall architecture of TRPM4 and identified key regulatory sites, including a Ca²⁺-binding pocket within the S1–S4 domain and an inhibitory ATP-binding site^9–12^. More recently, high-resolution structures expanded this conformational landscape by capturing apo closed, Ca²⁺-bound desensitized, Ca²⁺–PtdIns(4,5)P₂-bound open and ATP-inhibited states, thereby providing a structural framework for channel activation, desensitization and inhibition^13–15^.

An important advance came from the finding that TRPM4 conformational state and ligand recognition depend strongly on temperature. Structural analysis at physiological temperature revealed a temperature-dependent “warm” conformation of TRPM4 and showed that ligands can bind to different sites at 37 °C than at lower temperatures commonly used for cryo-EM sample preparation, indicating that ligand recognition in TRPM4 reflects the conformational ensemble available under a given set of environmental conditions rather than a single static structure^15^. In parallel, there has been growing interest in studying membrane proteins in systems that better preserve native lipid interactions^16–18^. Such approaches can improve the structural interpretation of membrane proteins by retaining lipid-associated features that influence conformational stability, gating and ligand recognition, particularly within transmembrane regulatory regions. Cell-derived membrane vesicles offer an attractive alternative to conventional detergent extraction because they retain endogenous membrane context and can preserve weakly associated lipids and cofactors that are often lost during solubilization^19,20^. Compared with reconstituted lipid nanodiscs or polymer-extracted native lipid particles, cell-derived vesicles can preserve larger membrane patches with a native-like lipid composition and local protein context. This may facilitate the identification of endogenous interaction partners and conformational states that could be lost or altered during detergent extraction, reconstitution or polymer solubilization^20,21^.

For TRPM4, previous structures obtained in detergent and nanodisc-like systems revealed cholesterol or cholesterol-like densities associated with the transmembrane domain, suggesting a defined paralipid environment, referring to lipids or lipid-like molecules that remain associated with the channel^10,13,22^. Anthranilic anilide derivatives have emerged as the best-characterised and most potent class of TRPM4 inhibitors, and structural work identified the binding site of NBA (4-chloro-2-(2-(naphthalene-1-yloxy)acetamido)benzoic acid) and IBA (4-chloro-2-(2-(3-iodophenoxy)acetamido)benzoic acid) in a pocket formed by the S3 and S4 helices, the S4–S5 linker and the TRP helix, a conserved cytosolic helix located C-terminal to S6, thereby establishing a structural basis for inhibitor recognition in the transmembrane domain^22^.

Although this previous study used styrene maleic acid (SMA) to extract TRPM4 and preserve aspects of its native paralipid environment, the structures were determined at 8 °C and without added Ca²⁺, conditions that are unlikely to favour an activated TRPM4 conformation. In parallel to this structural study, Gerber *et al*. performed a focused structure–activity relationship study of anthranilic anilide-based TRPM4 inhibitors by systematically modifying the phenoxy ring of the 4-chloro-2-(2-phenoxyacetamido) benzoic acid scaffold. This work identified PBA (4-chloro-2-(2-(3-(prop-2-yn-1-yloxy)phenoxy)acetamido) benzoic acid) as a more potent analogue relative to NBA, CBA (4-chloro-2-(2-(2-chlorophenoxy)acetamido)benzoic acid) and IBA, with improved ligand efficiency, aqueous solubility and reduced cytotoxicity compared to NBA^23^. However, the extent to which detergent-solubilised preparations reproduce the native lipid-associated architecture of TRPM4, and whether PBA engages the previously described inhibitor pocket, remains unknown. This information could be powerful for understanding the basis for continued improvement in small-molecule drug design targeting TRPM4. Here we combine cryo-EM with membrane vesicle preparations and temperature-controlled structural analysis to investigate TRPM4 ligand recognition under more physiological conditions. We first establish a vesicle-based structural workflow for TRPM4 and compare its lipid-associated architecture with detergent-solubilised TRPM4. We then determine the structures of TRPM4 in complex with PBA at low and physiological temperatures, and we find that PBA occupies distinct binding pockets depending on temperature. These findings reveal unexpected plasticity in TRPM4 ligand recognition and identify temperature as a key determinant of inhibitor binding within the TRPM4 transmembrane regulatory domain.

## Results

### Preparation of TRPM4-containing whole-cell vesicles

To examine TRPM4 in a membrane environment as close as possible to its native context, we established a workflow for the preparation of TRPM4-containing whole-cell vesicles from suspension HEK cells expressing C-terminally FLAG-tagged TRPM4, adapting the membrane vesicle strategy described by Tao *et al.*^19^. As outlined in Figure 1A, sonication generated membrane vesicles of mixed orientation, which were subsequently enriched by DEAE-Sepharose chromatography and purified by anti-FLAG affinity pull down, yielding inside-out vesicles containing TRPM4. Purified samples showed clear enrichment of TRPM4 on Coomassie-stained SDS–PAGE (Figure 1B). Cryo-EM micrographs revealed vesicles densely populated with TRPM4 particles, although vesicles with lower particle content were also observed (Figure 1C). Two-dimensional classification yielded well-defined class averages corresponding to both side and top views of TRPM4 in vesicles (Figure 1C).

**Figure 1:**
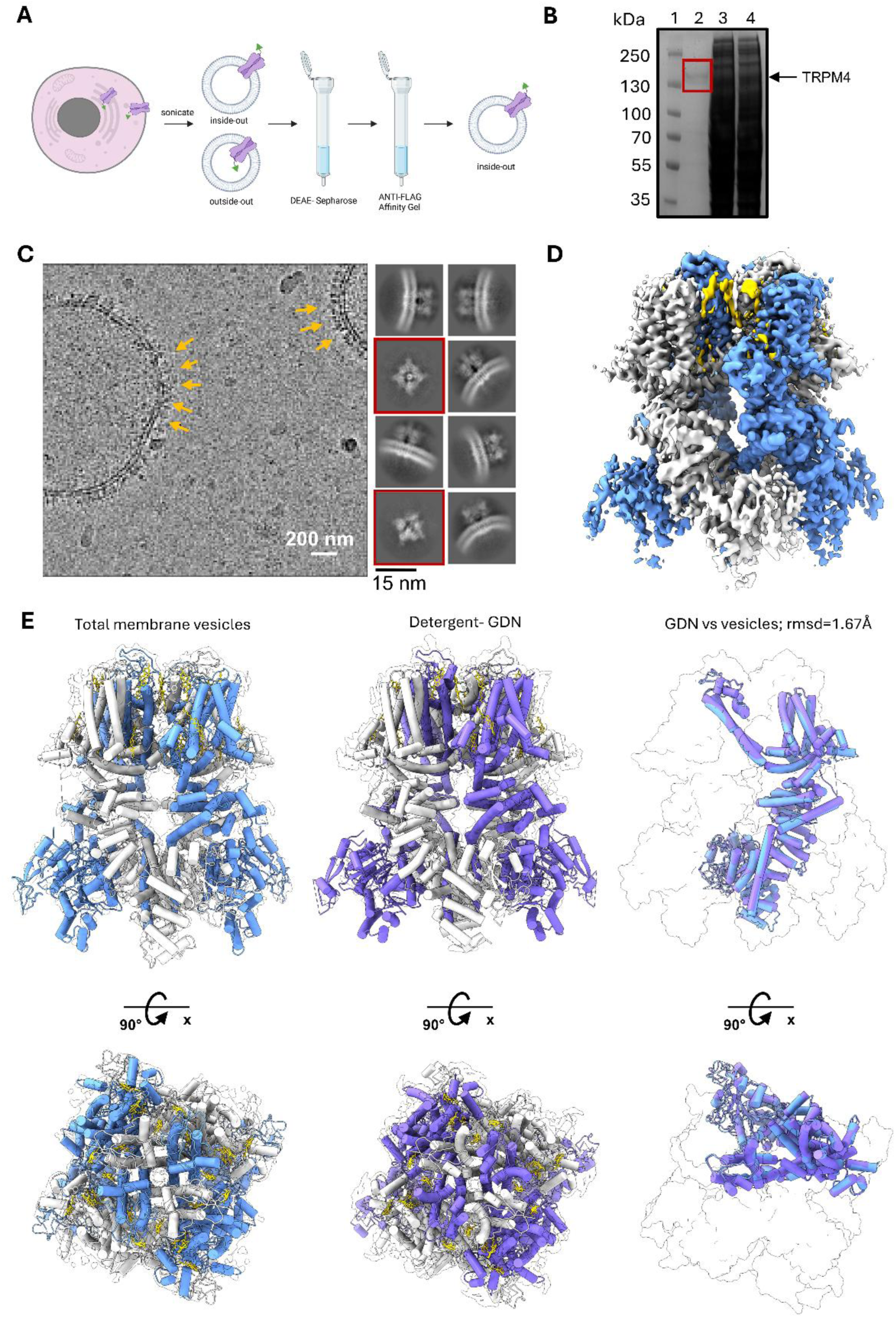
Purification, cryo-EM analysis and structural comparison of TRPM4 in whole-cell vesicles and GDN. **(A)** Schematic workflow for the generation and purification of TRPM4-containing whole-cell vesicles. The cell is shown in pink, TRPM4 in violet, and the intracellular FLAG tag as a green triangle. **(B)** Coomassie-stained SDS–PAGE gel showing TRPM4 enrichment during purification. Lane 1, molecular weight marker; lane 2, FLAG affinity-purified sample; lane 3, AEX-purified sample; lane 4, total cell lysate. The position of TRPM4 is indicated. **(C)** Representative cryo-EM micrograph of purified vesicles. Yellow arrows indicate TRPM4 densities embedded within the vesicle membrane. Representative 2D class averages derived from vesicle particles are shown on the right, including both side views and top views (red boxes). **(D)** Overall cryo-EM reconstruction of TRPM4 in whole-cell vesicles. The cryo-EM density is shown in white, and alternating subunits are highlighted in blue. Putative lipid-like densities associated with the transmembrane domain are highlighted in yellow. **(E)** Structural comparison of TRPM4 determined in whole-cell vesicles and in GDN. Left, TRPM4 structure obtained in vesicles with modelled cholesterol molecules shown in yellow and alternating subunits highlighted in blue. Middle, TRPM4 structure obtained in GDN with modelled cholesterol molecules shown in yellow and alternating subunits highlighted in violet. Right, superposition of the vesicle-derived and GDN-derived TRPM4 structures (RMSD = 1.67 Å).

Cryo-EM analysis of vesicle-embedded TRPM4 yielded a reconstruction at an overall resolution of 3.65 Å, with local resolution in the transmembrane domain reaching approximately 3.0 Å (Supplementary Figure 1, Table 1). The resulting map resolved the TRPM4 structure together with multiple lipid-like densities associated with the transmembrane domain, allowing visualization of the channel together with lipids retained from the native membrane environment (Figure 1D). For comparison, we also determined the structure of TRPM4 purified in GDN (glyco-diosgenin), which yielded a reconstruction with an overall resolution of 3.11 Å (Supplementary Figure 3, Table 1). Superposition of the vesicle-derived and GDN-derived structures revealed a high degree of structural similarity, with an RMSD of 1.67 Å (Figure 1E), indicating that detergent-solubilized TRPM4 in GDN closely resembles the architecture observed in whole-cell vesicles. By contrast, comparison of the vesicle-derived structure with previously reported TRPM4 structures revealed substantially greater differences for a nanodisc-derived structure (PDB 6BQR; RMSD = 4.081 Å), whereas a structure obtained in LMNG detergent (PDB 9MTA) remained highly similar (RMSD = 1.895 Å) (Supplementary Figure 2A, B)^10,13^. These observations suggest that the structural state sampled by TRPM4 may depend on the membrane-mimetic environment used for purification and structure determination. Together, these data establish whole-cell vesicles as a suitable system for structural analysis of TRPM4 and provide a basis for direct comparison with detergent-solubilized TRPM4.

**Table 1:**
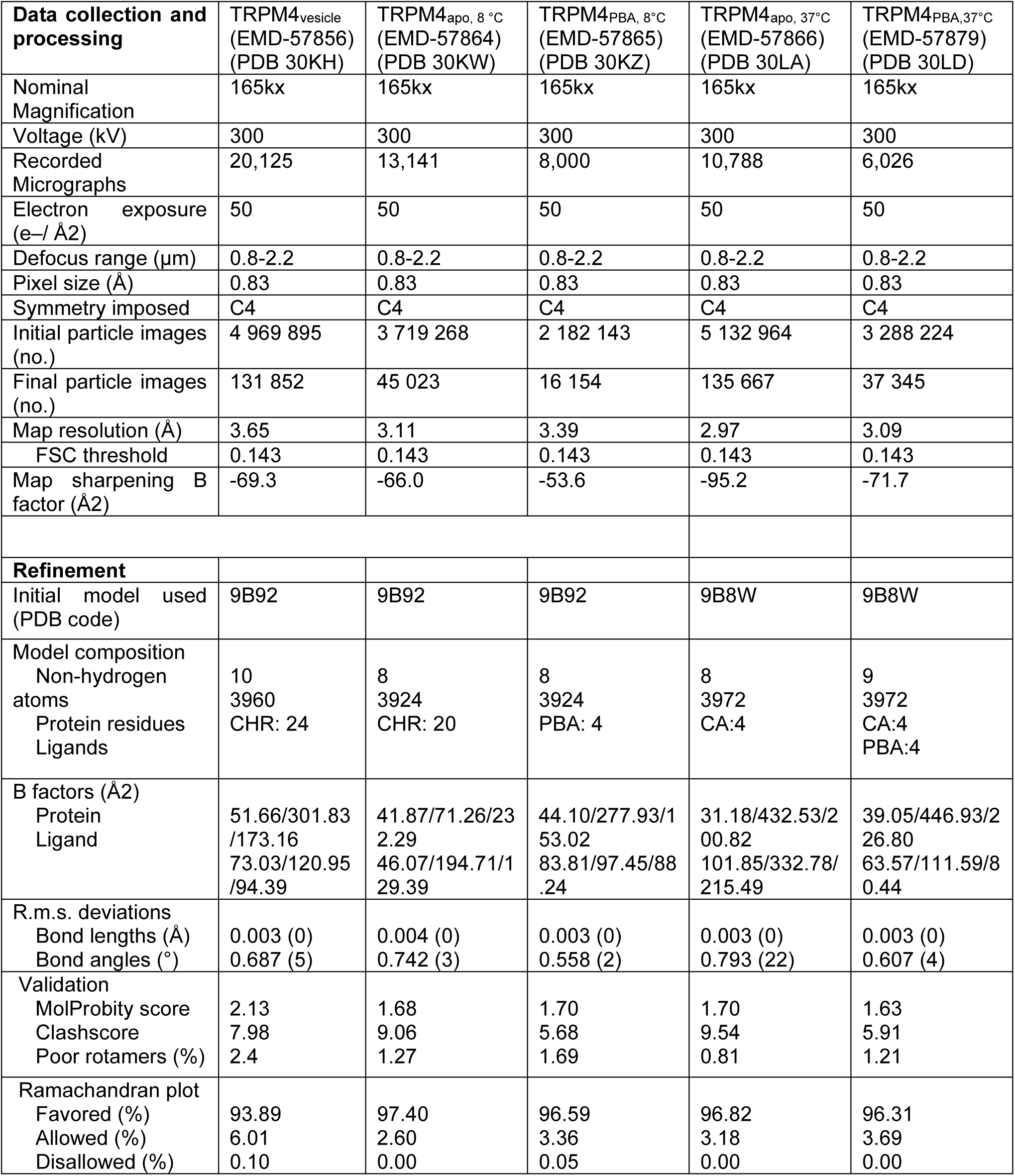
Cryo-EM data collection, refinement, and validation statistics for TRPM4 samples.

### TRPM4 in vesicles and GDN displays a similar cholesterol landscape

We next compared the lipid-associated architecture of TRPM4 in whole-cell vesicles and in GDN. In the vesicle-derived structure obtained at 8 °C, six cholesterol-like paralipid densities, modelled as cholesterol, could be resolved per subunit within the transmembrane region (Figure 2A–C). These densities were distributed across both membrane leaflets, with three cholesterol molecules assigned to the outer leaflet and three to the inner leaflet (Figure 2B, C). Although all six sites could be assigned, the densities corresponding to CHR-4, CHR-5 and CHR-6 in the inner leaflet were weaker than those of the remaining sites, suggesting lower occupancy.

**Figure 2:**
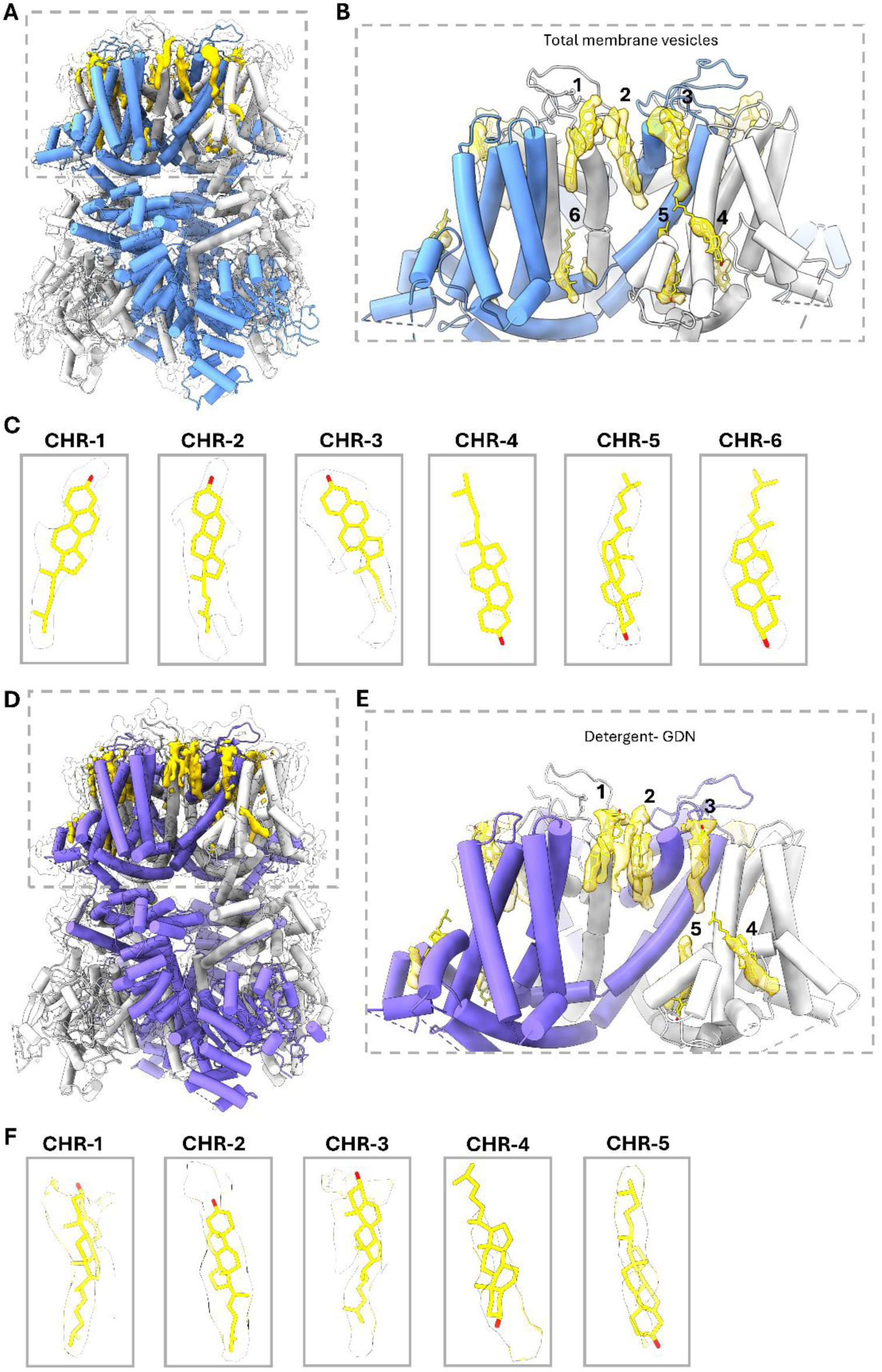
Cholesterol-binding sites in TRPM4 purified from whole-cell vesicles and GDN. **(A)** Overall structure of TRPM4 purified from whole-cell vesicles shown in cartoon representation, with alternating subunits coloured in light and dark blue. Cholesterol molecules modelled into lipid-like densities are shown in yellow. **(B)** Enlarged view of the transmembrane domain of vesicle-derived TRPM4 highlighting six cholesterol-binding sites per subunit, designated CHR-1 to CHR-6. **(C)** Individual views of cholesterol molecules CHR-1 to CHR-6 together with the corresponding cryo-EM densities obtained from the vesicle-derived structure. All densities are shown at the same contour level. **(D)** Overall structure of TRPM4 purified in GDN shown in cartoon representation, with alternating subunits coloured in light and dark violet. Cholesterol molecules modelled into lipid-like densities are shown in yellow. **(E)** Enlarged view of the transmembrane domain of GDN-purified TRPM4 highlighting five cholesterol-binding sites per subunit, designated CHR-1 to CHR-5. These sites correspond to those observed in the vesicle-derived structure, whereas CHR-6 was not resolved in GDN. **(F)** Individual views of cholesterol molecules CHR-1 to CHR-5 together with the corresponding cryo-EM densities obtained from the GDN-derived structure. All densities are shown at the same contour level.

To assess whether detergent extraction altered this lipid landscape, we determined the structure of TRPM4 in GDN at the same temperature (8 °C). In the GDN structure, five cholesterol molecules per subunit were resolved (Figure 2D–F). Importantly, these five sites corresponded to the same positions observed in the vesicle preparation, whereas one site, CHR-6, was only resolved in vesicles (Figure 2C, F). In our previous study, in which TRPM4 was extracted using SMA polymer, a cholesterol density at the corresponding position was likewise observed only at low occupancy, whereas this site was more prominently occupied in TRPM4 purified in LMNG/CHS detergent^22^. Thus, the cholesterol-binding pattern in GDN closely matched that observed in the vesicle-derived structure, indicating that the overall lipid organization surrounding TRPM4 is largely preserved following solubilization in GDN. The observed cholesterol arrangement is also consistent with our previous SMA-derived structure, suggesting that these lipid-binding sites represent stable features of the TRPM4 transmembrane domain^22^.

Furthermore, GDN has previously been used to investigate temperature– and ligand-dependent conformational changes in TRPM4^15^. Together with the close structural agreement between the vesicle-derived and GDN-derived structures (Figure 1E) and the highly similar cholesterol landscapes observed here (Figure 2), these findings indicate that the detergent-solubilized preparation preserves key features of the TRPM4 paralipid environment. Because vesicle samples prepared at 37 °C were not suitable for further structural analysis due to poorly defined densities in the 2D class averages, possibly reflecting increased mobility of TRPM4 within the vesicles or the presence of multiple conformational states that increased sample heterogeneity (Supplementary Figure 2C), whereas GDN samples remained amenable to high-resolution structure determination at 37 °C, we used GDN for subsequent analysis of PBA binding at low and physiological temperatures (Supplementary Figure 5, Table 1).

### PBA binds to distinct sites in TRPM4 at 8 °C and 37 °C

Having shown that TRPM4 displays a similar paralipid arrangement in GDN compared to in a native membrane environment, we next determined TRPM4 structures in complex with the potent anthranilic anilide inhibitor PBA at 8 °C, presenting a closed inactive state and at 37 °C in the presence of Ca^2+^, presenting a Ca^2+^-bound state (Supplementary Figures 4 and 6, Table 1). We also determined additional TRPM4 structures at 37 °C in the presence of Ca²⁺ but without PBA, and at 8 °C in the absence of PBA, providing ligand-free reference structures for comparison. At 8 °C, PBA was resolved in a pocket adjacent to S3 and S4 and bordered by the S4–S5 linker and the TRP helix (Figure 3A, D). This position corresponds to the inhibitor-binding site described for the related anthranilic anilides NBA and IBA in our previous study^22^. By contrast, in the 37 °C structure with Ca^2+^ ions present, PBA occupied a distinct site within the S1–S4 domain, close to the bound Ca²⁺ ion (Figure 3C, E). Thus, PBA shows a clear temperature-dependent relocation between two structurally distinct pockets.

**Figure 3:**
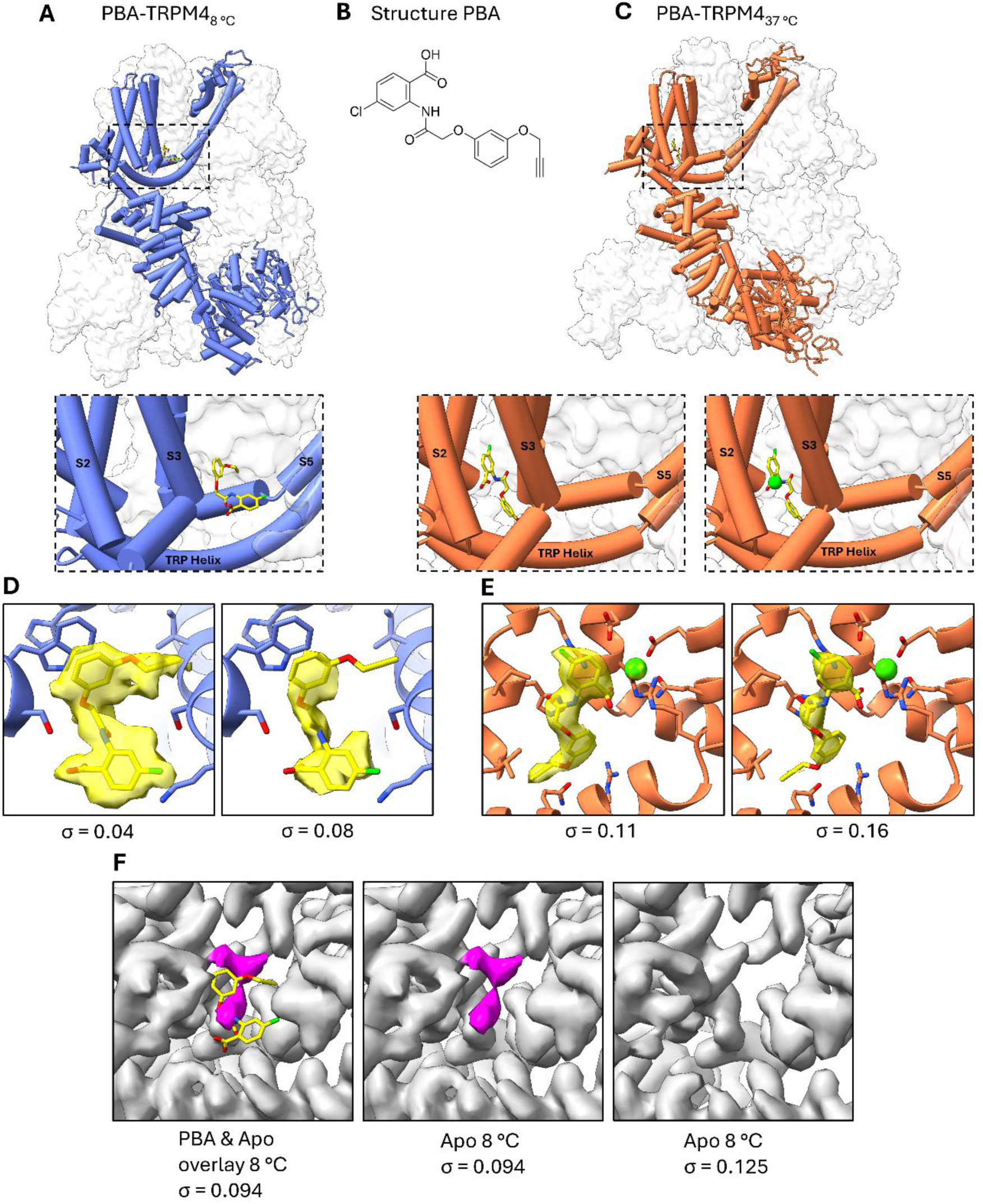
PBA binds to distinct sites in TRPM4 at 8 °C and 37 °C. **(A)** Structure of PBA-bound TRPM4_8 °C_ shown in cartoon representation, with the cryo-EM density shown as a transparent surface. The boxed region indicates the PBA-binding site located between S3, S4, S5 and the TRP helix. The lower panel shows an enlarged view of the binding pocket. **(B)** Chemical structure of PBA. **(C)** Structure of PBA-bound TRPM4_37 °C_ shown in cartoon representation, with the cryo-EM density shown as a transparent surface. The boxed region indicates the PBA-binding site within the S1–S4 domain, proximal to the bound Ca²⁺ ion (green sphere). The lower panel shows an enlarged view of the binding pocket. **(D,E)** Zoomed-in views of the PBA-binding sites in TRPM4 at 8 °C **(D)** and 37 °C **(E)**. PBA is shown with the corresponding cryo-EM density in yellow surface representation at two different density threshold levels, indicated by σ. **(F)** Comparison of the 8 °C PBA-bound structure with the corresponding region of the apo TRPM4 map. Left, overlay of PBA with a non-protein density present in the apo structure (highlighted in purple). Middle, the same region in the apo map at the same density threshold, showing no overlap with the PBA density. Right, the apo map is shown at a higher density threshold, where the purple density is no longer visible.

The density for PBA was well-defined in both structures and remained visible across different contour levels (Figure 3D, E), supporting the assignment of the ligand to each site. Additional inspection of the apo and PBA-bound maps confirmed the presence of ligand density only in the PBA-bound structures and revealed well-defined density for residues lining both binding pockets (Supplementary Figure 7). At 8 °C, the ligand density occupied the previously described transmembrane pocket, whereas at 37 °C the density shifted upward by approximately 12 Å into the S1–S4 region near the Ca²⁺-binding site. To exclude the possibility that the 8 °C PBA density had been misassigned from a pre-existing lipid-like density in the apo structure, we compared the ligand-bound and apo maps in the same region (Figure 3F, Supplementary Figure 7). In the apo map, an additional non-protein density is present near the 8 °C pocket and was highlighted in purple. However, this density does not overlap with the PBA density observed in the ligand-bound structure and is absent when PBA is present, arguing against misinterpretation of this apo density as bound PBA. Moreover, this apo density disappears at higher contour levels, consistent with comparatively low occupancy. Together, these observations support the assignment of PBA in the 8 °C structure and indicate that the apo density likely represents a weakly occupied lipid-like species.

### Temperature-dependent PBA binding is accompanied by a shift in pocket interactions

To define the structural basis of the two PBA-binding modes, we analysed the residues surrounding the ligand in both structures (Figure 4). In the 8 °C structure, PBA is coordinated within the previously described inhibitor pocket by residues lining the interface between the S3 helix, S4 helix, S4–S5 linker and TRP helix. Ser863 on the S3 helix is positioned near the phenoxyacetamide linker of PBA, whereas Ser924 and Lys925 in the S4–S5 linker interact with the chlorinated anthranilic acid ring. The aromatic side chains of Trp864 on the S3 helix and His908 on the S4 helix form a triple π-stacking arrangement with the phenoxy ring of PBA. Additional hydrophobic contacts are provided by Ile920 in the S4–S5 linker, which packs against the propargyloxy group, and by Tyr1057 in the TRP helix, which contacts the anthranilic acid moiety (Figure 4A).

**Figure 4:**
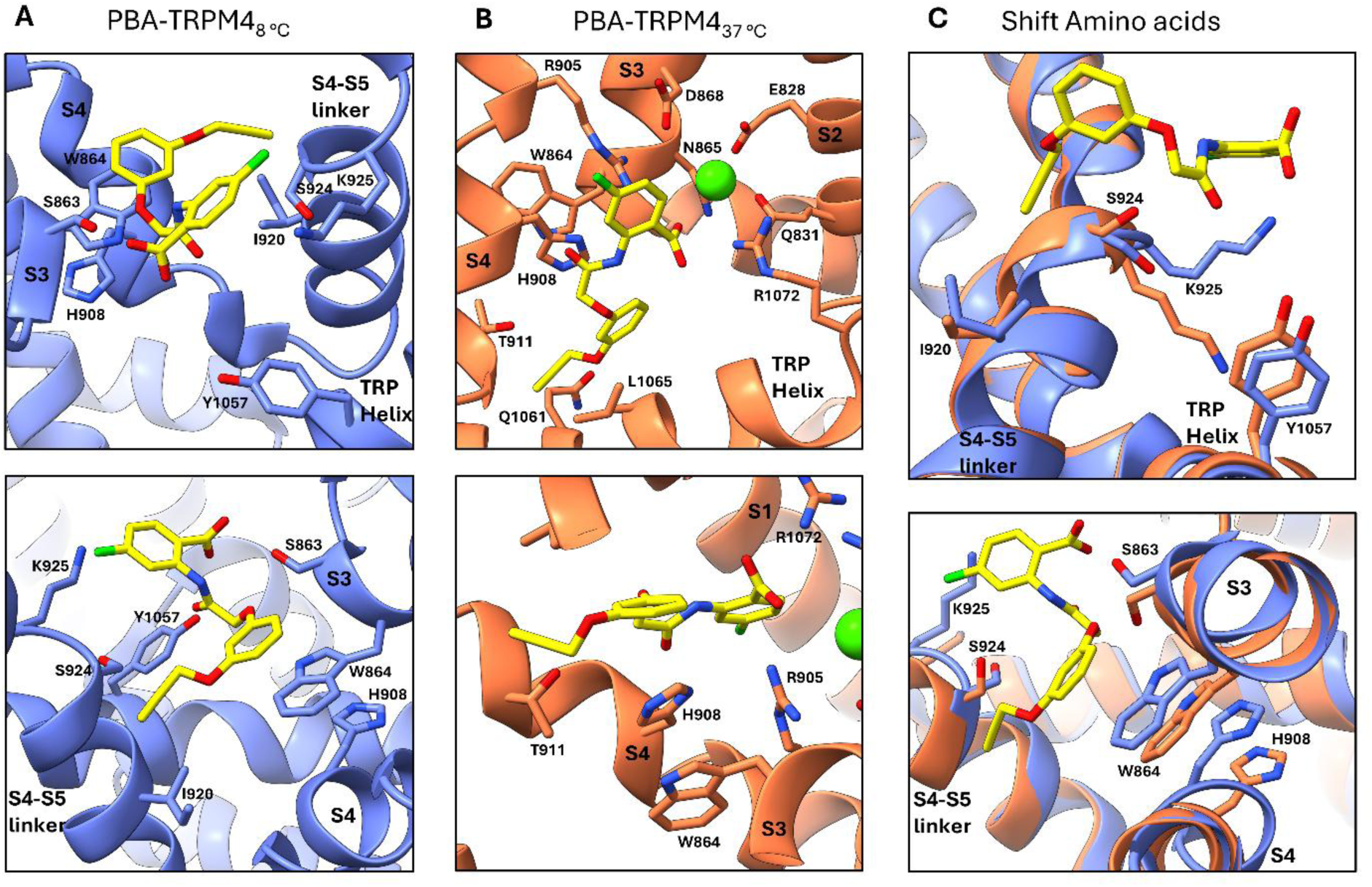
Temperature-dependent interactions of PBA within the TRPM4 binding pocket. **(A)** Amino acid residues of TRPM4 involved in PBA binding at 8 °C. The lower panel highlights the triple π-stacking interaction of PBA with W864 and H908. **(B)** Amino acid residues of TRPM4 involved in PBA binding at 37 °C. The lower panel highlights the corresponding triple π-stacking interaction of PBA with W864 and H908, with the ligand engaging the opposite side of these residues compared to 8 °C. **(C)** Overlay of the TRPM4 structures at 8 °C and 37 °C showing the shift in PBA position and the associated rearrangement of interacting residues. The lower panel highlights the corresponding shift in the residues involved in the π-stacking interaction.

In the 37 °C structure, PBA is surrounded by a different set of residues within the S1–S4 domain (Figure 4B). The ligand is positioned close to the Ca²⁺-binding region, where Glu828 and Gln831 on the S2 helix, together with Asn865 and Asp868 on the S3 helix, coordinate the bound Ca²⁺ ion. In this binding mode, Arg905 on the S4 helix contacts the anthranilic acid moiety of PBA, whereas Thr911 on the S4 helix and Gln1061 on the TRP helix are positioned near the propargyloxy group. Arg1072 on the TRP helix interacts with the chlorinated anthranilic acid ring, while Leu1065 on the TRP helix provides hydrophobic contacts with the phenoxy ring and propargyloxy group. Despite the relocation of the ligand, the triple π-stacking interaction with the phenoxy ring is retained, again involving Trp864 on the S3 helix and His908 on the S4 helix. Notably, however, PBA engages the opposite side of these residues compared with the 8 °C structure, indicating a different local arrangement of the aromatic core relative to the π-stacking pair.

Direct comparison of the two structures revealed a coordinated rearrangement of the binding pocket accompanying ligand relocation (Figure 4C). Trp864 and His908 undergo a pronounced positional shift while maintaining the π-stacking interaction with PBA. In parallel, Ser863 and Ser924 rotate away from the ligand in the 37 °C structure, whereas Ile920 and Lys925 also adopt altered positions relative to the 8 °C pocket. Together, these changes reshape the local environment of the ligand and support the view that temperature-dependent conformational differences in the transmembrane domain enable two distinct PBA-binding modes.

### A structural model for temperature-dependent PBA recognition in TRPM4

Based on these structures, we propose a model in which PBA recognition by TRPM4 is governed by temperature-dependent changes in pocket accessibility and local residue arrangement (Figure 5). In the apo state at 8 °C, TRPM4 adopts a closed conformation. In the presence of Ca²⁺ and PBA, the channel retains a non-conducting pore architecture at both temperatures, but the position of the ligand differs markedly. At 8 °C, PBA occupies the canonical inhibitor pocket adjacent to S3, S4, the S4–S5 linker and the TRP helix. At 37 °C, PBA relocates to a distinct site within the S1–S4 domain, proximal to the Ca²⁺-binding region.

**Figure 5:**
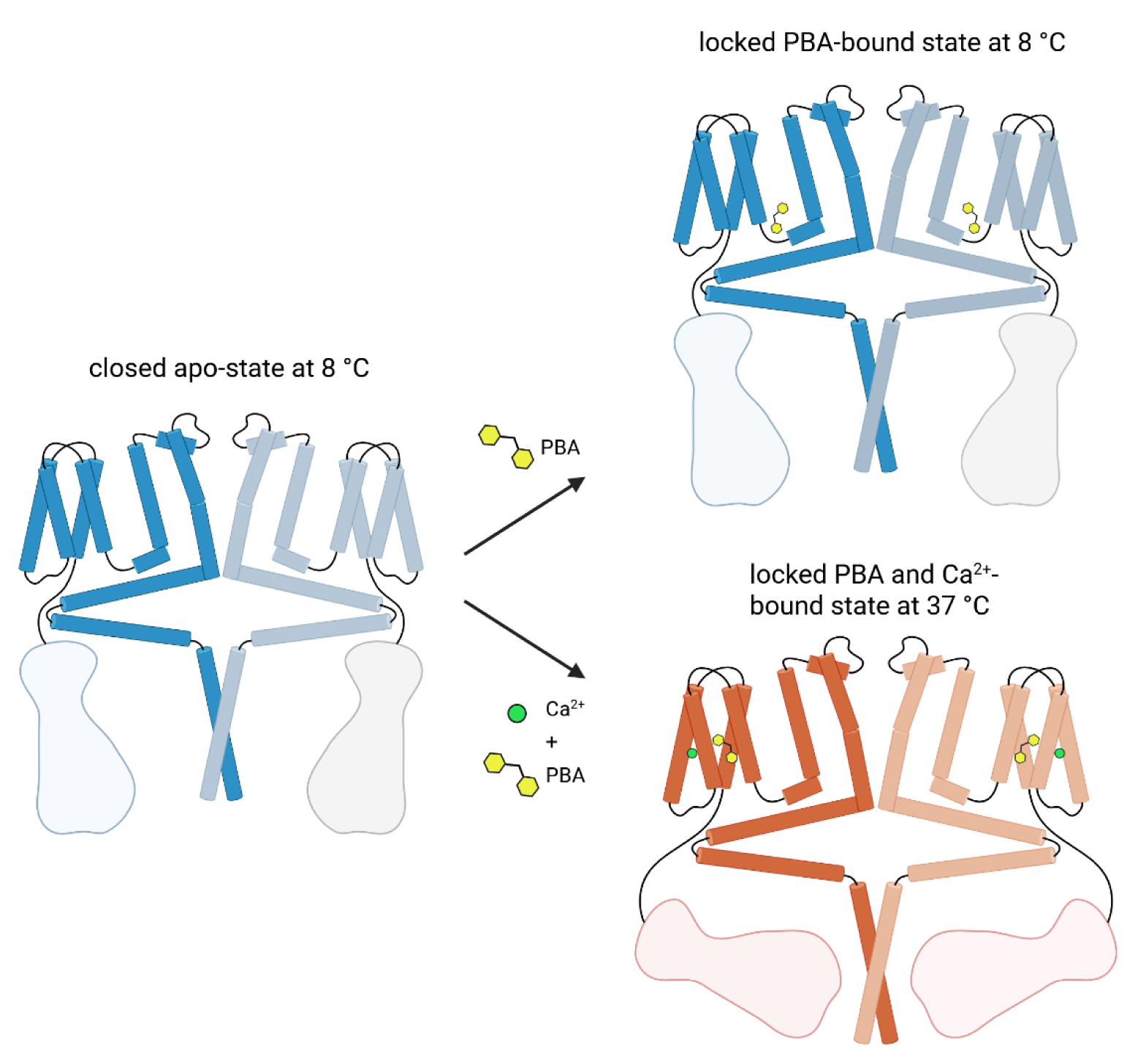
Schematic model of temperature-dependent PBA binding in TRPM4. Schematic representation of TRPM4 illustrating the distinct PBA-binding sites observed at 8 °C and 37 °C. In the apo state at 8 °C, TRPM4 is shown in the closed conformation. In the PBA-bound structures, PBA occupies different binding pockets under the two conditions analysed: at 8 °C without added Ca²⁺ and at 37 °C in the presence of Ca²⁺. At 8 °C, PBA binds in the previously described inhibitor pocket adjacent to S3, S4, the S4–S5 linker and the TRP helix, whereas at 37 °C it relocates to a distinct site within the S1–S4 domain proximal to the Ca²⁺-binding region.

This model provides a structural framework for accommodating a single inhibitor scaffold in two distinct pockets within the TRPM4 transmembrane region. More broadly, these findings identify the S1–S4 module as a structurally plastic region capable of supporting temperature-dependent ligand recognition, consistent with the emerging view that ligand binding in TRPM4 depends on environmental conditions and conformational state^13,15^.

## Discussion

Our study provides three main findings. First, whole-cell vesicles enabled structural analysis of TRPM4 in a membrane environment that preserves lipid-associated features of the channel (Figures 1 and 2). Second, comparison of vesicle-derived and GDN-solubilized TRPM4 revealed a highly similar lipid-binding landscape, indicating that GDN-solubilized TRPM4 retains key aspects of its paralipid environment under our conditions (Figure 2). Third, and most importantly, we find that the anthranilic anilide inhibitor PBA occupies distinct binding pockets at 8 °C and 37 °C, revealing temperature-dependent plasticity in ligand recognition within the TRPM4 transmembrane regulatory region (Figures 3–5). These two binding sites are also very likely to be used by the other anthranilic anilide inhibitors including NBA, CBA and IBA.

The vesicle-based part of this work was motivated by the concern that detergent extraction can disrupt weakly associated lipids and alter the structural environment of membrane proteins. Tao *et al*. showed that membrane vesicles can preserve native membrane context and support high-resolution cryo-EM analysis without prior detergent extraction^19^. In our study, TRPM4 structures obtained from whole-cell vesicles and GDN were highly similar in the transmembrane region, particularly regarding cholesterol occupancy (Figures 1 and 2). Consistent with this observation, the vesicle-derived structure displayed a substantially lower RMSD when compared with a TRPM4 structure obtained in LMNG detergent than with a previously reported nanodisc-derived structure (Supplementary Figure 2A, B). Five cholesterol-binding sites were shared between the two preparations, whereas a sixth site was only resolved in vesicles and displayed weaker density, consistent with lower occupancy (Figures 2B,D). These data indicate that GDN-solubilized TRPM4 can preserve the major cholesterol-binding pattern relevant for structural analysis, while vesicles remain useful for defining native-like lipid interactions more directly. The most unexpected finding of this study is that PBA occupies two different binding sites depending on temperature. At 8 °C, PBA binds in a pocket formed by S3, S4, the S4–S5 linker and the TRP helix, in agreement with the binding site previously identified for NBA and IBA (Figures 3A, D)^22^. At 37 °C, however, PBA relocates into the S1–S4 domain, close to the Ca²⁺-binding region (Figures 3C, E). This shift is accompanied by a coordinated reorganization of surrounding residues while retaining the triple π-stacking interaction with Trp864 and His908 (Figure 4). These observations indicate that the TRPM4 transmembrane domain can support multiple binding modes for a single inhibitor scaffold, and that temperature can shift the balance between them.

This conclusion is consistent with the emerging view that temperature affects the TRPM4 conformational landscape and ligand recognition. Hu et al. showed that TRPM4 adopts a temperature-dependent warm conformation at physiological temperature and that ligands can bind to different sites at 37 °C than at lower temperatures^15^. Our data extend this principle by showing temperature-dependent ligand relocation, in which PBA relocates between two pockets under otherwise comparable biochemical conditions (Figure 3). The 37 °C PBA site is also notable because of its proximity to the Ca²⁺-binding pocket in the S1–S4 domain. Recent structural work identified this region as a central regulatory hub in TRPM4 gating, with Ca²⁺, PtdIns(4,5)P₂ and ATP coupling through the S1–S4 domain, S4–S5 linker and TRP helices to control pore opening, desensitization and inhibition^13^. Earlier functional work showing that PtdIns(4,5)P₂ rescues TRPM4 from desensitization further supports this view^24^. In our 37 °C structure, PBA occupies a pocket framed by residues that include Glu828, Gln831, Asn865 and Asp868 from the Ca²⁺-binding region, together with Arg905, Thr911, Gln1061, Leu1065 and Arg1072 (Figure 4B). Although our data do not define the functional consequence of this binding mode, they provide a structural basis for why PBA may differ from simpler anthranilic anilides in potency and recognition mode.

This point is especially relevant in the context of the PBA scaffold. PBA emerged from a focused SAR campaign as a more potent analogue than NBA, with improved ligand efficiency, aqueous solubility and lower cytotoxicity^23^. The 8 °C structure shows that PBA remains compatible with the canonical anthranilic anilide pocket, as expected from its close chemical relationship to NBA, whereas the 37 °C structure reveals an additional recognition mode that would not have been predicted from low-temperature structures alone (Figure 3). The ability of PBA to occupy more than one binding site depending on temperature, as shown here, may help explain its high potency relative to other known inhibitors. An important control in this study is the comparison of the 8 °C PBA-bound map with the corresponding apo density (Figure 3F). In the apo structure, a weak non-protein density is present near the canonical pocket, but it does not overlap with the PBA density and disappears at higher contour levels, consistent with low occupancy. This argues against ligand misassignment and instead suggests that the apo site may occasionally be occupied by a weakly bound lipid-like species. We therefore favour the interpretation that PBA does not simply bind more tightly within the canonical pocket, but that its scaffold is compatible with multiple temperature-dependent recognition states in TRPM4.

Our study has limitations. We do not assign functional consequences to the two PBA-binding modes, nor do we determine whether the 37 °C site can also be sampled by other anthranilic anilides under appropriate conditions. In addition, vesicle-derived structures at 37 °C could not be obtained because of insufficient data quality (Supplementary Figure 2C), so the physiological-temperature analysis relies on GDN-solubilized material (Figure 2). Nonetheless, the strong overlap between vesicle and GDN cholesterol landscapes at 8 °C supports the structural relevance of the detergent-solubilized preparation for this comparison. Furthermore, GDN-solubilized TRPM4 has previously been used to investigate temperature– and ligand-dependent conformational changes in TRPM4^15^. More broadly, these findings fit into a longer trajectory of TRPM4 research showing that the channel is regulated by intracellular Ca²⁺, PtdIns(4,5)P₂, ATP and other cellular factors, and that altered TRPM4 function is relevant to human disease, including cardiac conduction disorders^2,25^. Our structures now add temperature-dependent inhibitor recognition to this framework.

An important next step will be to determine whether temperature-dependent site switching is unique to PBA or represents a broader property of anthranilic anilides and related TRPM4 ligands. More generally, our work suggests that structurally informed drug discovery for TRPM4 may benefit from explicitly incorporating physiological variables into screening and structure determination. In that sense, the combination of vesicle-based structural analysis with temperature-controlled cryo-EM provides a useful framework for studying environment-dependent pharmacology in TRPM4 and potentially in other temperature-sensitive ion channels.

## Methods

### Design and generation of plasmid constructs

For this study, the codon-optimized wild-type *Homo sapiens* TRPM4 gene encoding the full-length protein was synthesized, equipped with consecutive C-terminal FLAG tags, and inserted into the pCDNA3.1 vector under control of the CMV promoter for expression in HEK293 cells (GenScript Biotech).

### Expression and purification of TRPM4

Full-length human TRPM4 was expressed and purified from suspension-cultured HEK293F cells. For transient expression, cells were transfected with 1.5 mg of the TRPM4 plasmid per liter of culture using polyethylenimine (PEI). Cultures were maintained at 37 °C and 5 % CO₂ for 48 h before harvesting by centrifugation at 3,000 × g for 30 min at 4 °C. Pellets were washed once with PBS and collected by an additional centrifugation step.

For detergent solubilization, 20 g of cell pellets were resuspended in lysis buffer (25 mM HEPES-NaOH pH 7.4, 200 mM NaCl) supplemented with cOmplete™ EDTA-free protease inhibitors (Roche), two tablets per 100 mL were used. Cells were lysed by a single passage through an LM20 Digital Microfluidizer at 18,000 psi. Membranes were isolated by ultracentrifugation (Optima XPN-100, Ti45 rotor, 30,000 rpm, 30 min, 4 °C) and resuspended in 40 mL solubilization buffer (25 mM HEPES-NaOH pH 7.4, 200 mM NaCl, 1 % GDN) containing one tablet of protease inhibitor. The membrane suspension was homogenized using a 100 mL Dounce homogenizer (Sigma) and incubated under gentle stirring for 3 h at 4 °C. Insoluble material was removed by ultracentrifugation (Optima XPN-100, Ti45 rotor, 30,000 rpm, 30 min, 4 °C), and the supernatant containing solubilized C-terminally FLAG-tagged TRPM4 was incubated with 1 mL FLAG® M2 affinity gel (Millipore) per 50 mL volume for 3 h at 4 °C under continuous stirring. After binding, the resin was transferred to an Econo-Pac® gravity-flow column (Bio-Rad), the resin was washed with 40 mL wash buffer (25 mM HEPES-NaOH pH 7.4, 200 mM NaCl, 0.02 % GDN), and the protein was eluted with 5 mL elution buffer (25 mM HEPES-NaOH pH 7.4, 200 mM NaCl, 0.02 % GDN, 120 µg/ mL 3×FLAG peptide). The eluate was concentrated to 500 µL using a 100 kDa Amicon Ultra-4 concentrator and further purified by size-exclusion chromatography on a Superose 6 column. Peak fractions were concentrated again (100 kDa Amicon) to an A₂₈₀ of 1.0 for cryo-EM grid preparation.

For preparation of total membrane vesicles, 20 g of cell pellets were resuspended in 100 mL solubilization buffer (25 mM HEPES-NaOH pH 7.4, 200 mM NaCl) supplemented with two tablets of cOmplete™ EDTA-free protease inhibitors. The suspension was homogenized with 30 strokes in a 100 mL Dounce grinder (Sigma), transferred to a 250 mL metal beaker, and sonicated using a Branson 250 Digital Sonifier (102-C converter, ½-inch tip) at 60 % amplitude for four cycles (30 s on / 30 s off). The lysate was clarified by centrifugation (12,000 × g, 10 min, 4 °C; Beckman Avanti J-20 XP, JA-20 rotor). The supernatant was applied to a 4 mL DEAE Sepharose gravity column equilibrated with solubilization buffer. The flowthrough was collected, and the resin was washed with three column volumes of lysis buffer. Flowthrough and wash fractions were pooled, distributed into 50 mL tubes, and incubated with 1 mL FLAG® M2 affinity gel per tube for 1 h at 4 °C under continuous rotation. After incubation, the suspension was transferred to an Econo-Pac® column, the resin was washed with 40 mL solubilization buffer and the protein was eluted with 5 mL elution buffer (25 mM HEPES-NaOH pH 7.4, 200 mM NaCl, 120 µg/ mL 3×FLAG peptide). The eluate was concentrated to ∼50 µL using a 100 kDa Amicon Ultra-4 concentrator for cryo-EM grid preparation.

### SDS-PAGE analysis

SDS–PAGE was used to assess protein purity throughout the purification workflow. Protein samples (37.5 µL) were mixed with 12.5 µL of 4× NuPAGE LDS Sample Buffer (Thermo Scientific) and heated at 95 °C for 5 min. Samples were loaded onto 4–12 % SurePAGE™ Bis-Tris precast gels (Witec AG), alongside a Spectra™ Prestained Protein Ladder (Thermo Scientific; 10–180 kDa). Electrophoresis was carried out in 1× Tris-MOPS SDS running buffer (Witec AG) at 200 V for 30 min. Gels were rinsed with Milli-Q water and stained for 2 h with QuickBlue Protein Stain (LuBioScience GmbH) under shaking. After washing in Milli-Q water, gels were imaged using an iBright FL1500 Imaging System (Thermo Scientific).

### Cryo-EM sample preparation and data collection

Purified TRPM4, either solubilized in GDN or retained in total membrane vesicles, was used directly for grid preparation or subjected to ligand-binding conditions with PBA. A 10 mM PBA stock solution was prepared in DMSO. For binding experiments, samples were adjusted to a final PBA concentration of 0.5 mM and incubated for 15 min at room temperature prior to vitrification. Unless stated otherwise, samples were maintained on ice before grid preparation. For temperature-dependent experiments at 37 °C, CaCl₂ was added to a final concentration of 5 mM, and samples were incubated at 37 °C for 10 min immediately prior to freezing. For cryo-EM, 400-mesh gold UltrAuFoil R1.2/ 1.3 grids (Quantifoil) were rendered hydrophilic by glow discharge (15 mA, 60 s) using a PELCO EasyGlow device (TED Pella). Then, 3 µL of concentrated protein solution was applied to the grids, and excess liquid was blotted for 2.5 s, followed by rapid vitrification in liquid ethane using a Vitrobot Mark IV (Thermo Fisher Scientific). For experiments performed at 8 °C or 37 °C, the Vitrobot chamber temperature was adjusted accordingly and maintained at 100 % relative humidity.

Cryo-EM data acquisition was performed using the automated EPU software on a Titan Krios G4 transmission electron microscope (Thermo Fisher Scientific) operated at 300 kV and equipped with a cold field emission gun and a Falcon4 direct electron detector. Micrographs were collected in counting mode at a nominal magnification of 165,000×, yielding a calibrated pixel size of 0.83 Å. Data were recorded over a defocus range of 0.8–2.2 µm with a total exposure of 50 e⁻/ Å², and stored in Electron Event Recording (EER) format.

### Cryo-EM data processing and model refinement

Cryo-EM image processing was performed using cryoSPARC (v5.0.2)^26^. Patch-based motion correction implemented in cryoSPARC was applied to align the movie stacks and to perform dose-weighting. CTF parameters were estimated using the patch-based method.

For the TRPM4_vesicle_ dataset, a total of 20,125 movies were collected at a pixel size of 0.83 Å. An initial set of 500 particles was manually picked and subjected to one round of 2D classification to generate templates. These templates were then used for automated particle picking across the dataset, resulting in 4,969,895 particles with a box size of 440 pixels. Two rounds of 2D classification were performed for particle cleaning, yielding 293,672 particles after the first round and 32,963 particles after the second round. Ab initio reconstruction followed by non-uniform refinement resulted in a single 3D map with an overall resolution of 3.39 Å in C4 symmetry (Supplementary Figure 1).

For the data of the TRPM4_apo, 8°C_, a total of 13,141 movies at 0.83 Å per pixel were collected. An initial set of 1000 particles was manually picked and subjected to one round of 2D classification to generate templates. Template-based automated particle picking resulted in a set of 3,719,268 particles at 560 pixels. One round of 2D classification was performed resulting in 128,167 particles. Ab initio reconstruction yielded three 3D reconstructions. One reconstruction representing 45,023 particles was selected for further non-uniform refinement, resulting in a map at 3.11 Å overall resolution in C4 symmetry (Supplementary Figure 3).

For the TRPM4_PBA, 8 °C_ dataset, a total of 8,000 movies were recorded at a pixel size of 0.83 Å. 2D class averages obtained from the HsTRPM4_apo_ dataset were used as templates for particle picking. Template-based automated picking yielded 2,182,143 particles extracted with a box size of 440 pixels. A single round of 2D classification was performed for particle cleaning, resulting in 175,909 particles. Ab initio reconstruction yielded three 3D reconstructions. One reconstruction containing 16,518 particles was selected for further non-uniform refinement, yielding a 3D map with an overall resolution of 3.39 Å under C4 symmetry (Supplementary Figure 4).

For the TRPM4_apo, 37 °C_ dataset, a total of 10,788 movies were recorded at a pixel size of 0.83 Å. 2D class averages from the HsTRPM4_apo_ dataset were used as templates for particle picking. Template-based automated picking resulted in 5,132,964 particles extracted with a box size of 560 pixels. A single round of 2D classification was carried out for particle cleaning, yielding 309,801 particles. Ab initio reconstruction yielded two classes, of which one contained 170,133 particles. These particles were subsequently subjected to ab initio reconstruction and non-uniform refinement, resulting in a single 3D map with an overall resolution of 2.97 Å under C4 symmetry (Supplementary Figure 5).

For the TRPM4_PBA, 37 °C_ dataset, a total of 6,026 movies were recorded at a pixel size of 0.83 Å. 2D class averages from the HsTRPM4_apo_ dataset were used as templates for particle picking. Template-based automated picking yielded 3,288,224 particles extracted with a box size of 440 pixels. Two rounds of 2D classification were performed for particle cleaning, resulting in 102,591 particles after the first round and 37,345 particles after the second round. These particles were subsequently used for ab initio reconstruction followed by non-uniform refinement, yielding a 3D map with an overall resolution of 3.09 Å under C4 symmetry (Supplementary Figure 6).

Atomic models for TRPM4_vesicle,_ TRPM4_apo, 8°C,_ TRPM4_PBA, 8 °C_, TRPM4_apo, 37 °C_ and TRPM4_PBA, 37 °C_ structures were mainly built in Coot 1.1.19^27^, using a model PDB id: 9B92 as an initial model for the 8 °C structures and 9B8W for the 37 °C Structures. Real-space refinement for all built models was performed using Phenix, version 1.21.2-5419-000, by applying a general restraints setup^28^.

### Data visualisation

Molecular graphics and analyses were carried out using UCSF ChimeraX (version 1.10), developed by the Resource for Biocomputing, Visualisation, and Informatics at the University of California, San Francisco, and supported by NIH grant R01-GM129325 and the Office of Cyber Infrastructure and Computational Biology (NIAID)^29^.

### Data Availability

The reconstructed maps are available from the EMDB database under access codes TRPM4_vesicle_ (EMD-57856), TRPM4_apo, 8°C_ (EMD-57864), TRPM4_PBA, 8 °C_ (EMD-57865), TRPM4_apo, 37 °C_ (EMD-57866) and TRPM4_PBA, 37 °C_ (EMD-57879). The atomic models are available in the PDB database, access codes TRPM4_vesicle_ (PDB 30KH), TRPM4_apo, 8°C_ (PDB 30KW), TRPM4_PBA, 8 °C_ (PDB 30KZ), TRPM4_apo, 37 °C_ (PDB 30LA) and TRPM4_PBA, 37 °C_ (PDB 30LD).

## Acknowledgements

Cryo-EM data were in part collected at the Dubochet Center for Imaging (DCI) in Lausanne. The DCI-Lausanne is a joint initiative of the EPFL and the Universities of Lausanne and Geneva. We are grateful to Bertrand Beckert, Emiko Uchikawa, Sergey Nazarov for their valuable assistance. We further acknowledge Kelvin Lau, Florence Pojer, Laurence Durrer and Soraya Quinch from the PTPSP at the EPFL for their support with protein expression. We also thank Christian Gerber and Martin Lochner for kindly providing PBA.

## Contributions

HA, HS, and BE conceptualised the project. DS and BE performed the cryo-EM sample preparation and structural determination, with support from the DCI, Lausanne. BE carried out project administration and supervision. HS and HA acquired funding. JSR contributed to project organisation and data interpretation. DS and BE wrote the first draft of the manuscript. All authors contributed to manuscript review and editing. All authors have read and approved the final version of the manuscript.

## Supplementary Figures

**Supplementary Figure 1:**
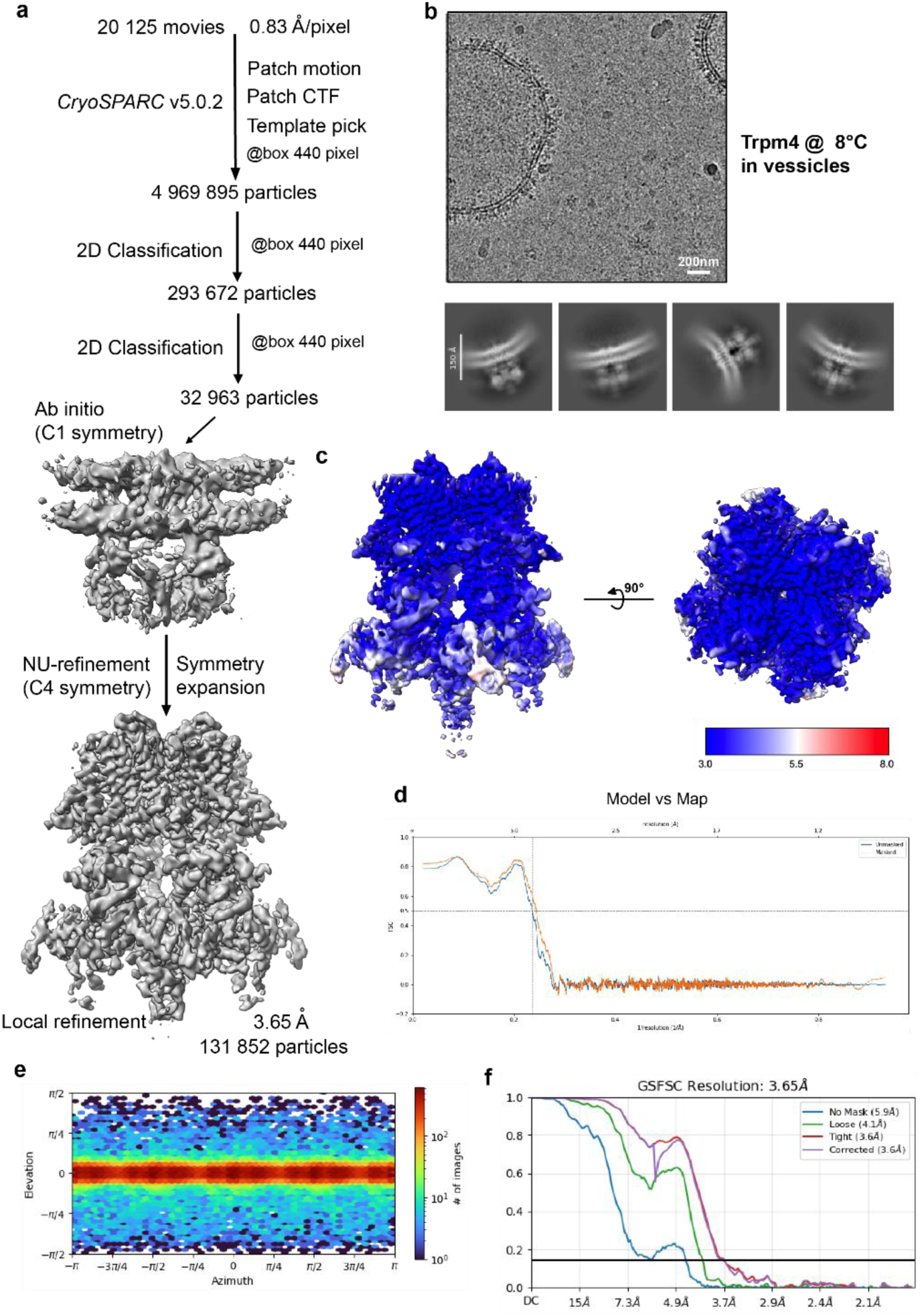
Cryo-EM data processing workflow and map resolution. (a) The image processing workflow of TRPM4_vesicle_. (b) micrograph and 2D classes (c) Local resolution (d) Model vs map FSC curves. (e) particle direction distribution. (f) FSC curves.

**Supplementary Figure 2:**
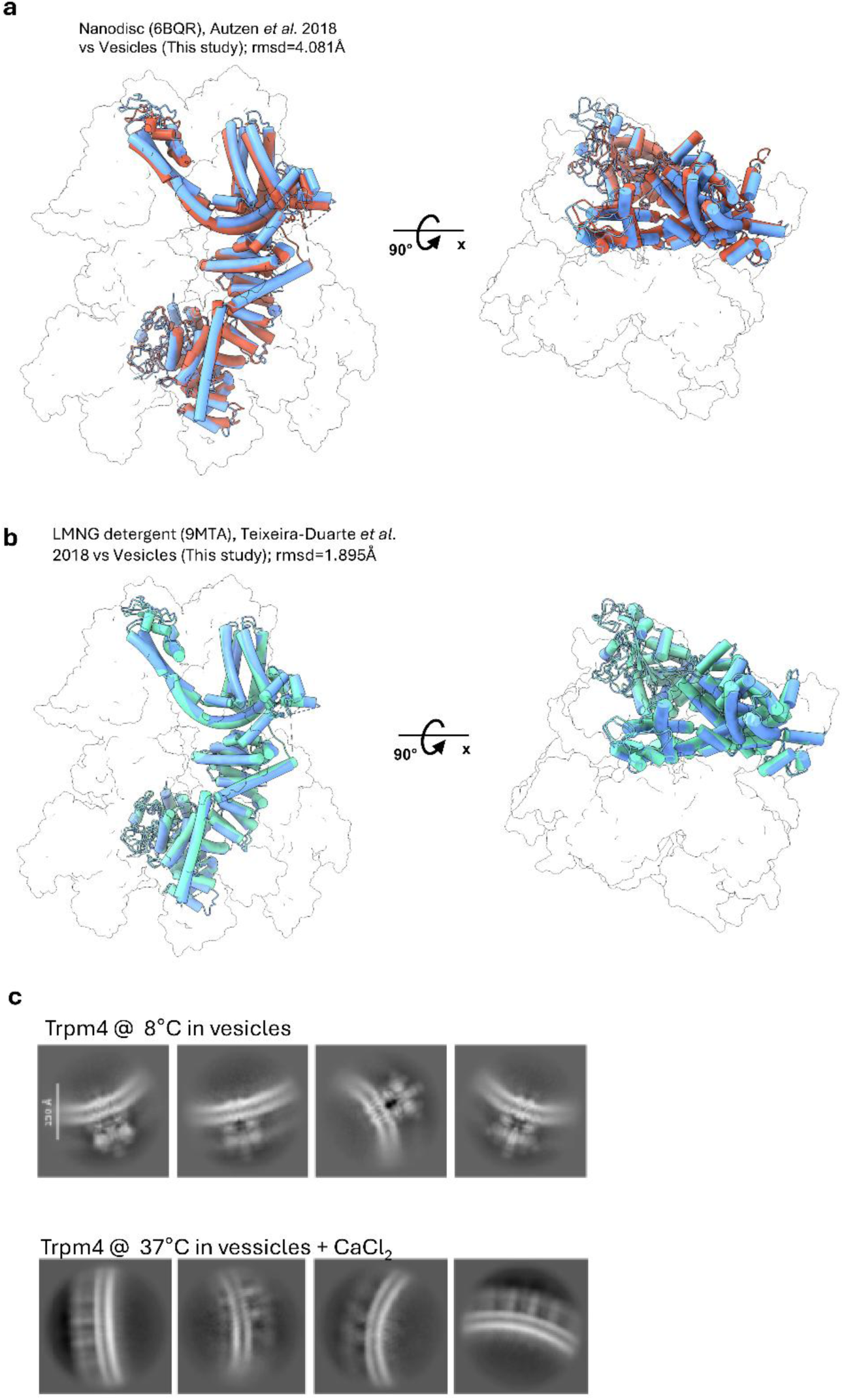
Structural comparison of vesicle-derived TRPM4 with previously reported structures and assessment of vesicle samples prepared at different temperatures. **(a)** Superposition of the TRPM4 structure determined in whole-cell vesicles in this study (blue) with the previously reported TRPM4 structure obtained in nanodiscs (PDB 6BQR; salmon) from Autzen *et al*^10^. Side and top views are shown. The two structures superimpose with a root mean square deviation (RMSD) of 4.081 Å. **(b)** Superposition of the TRPM4 structure determined in whole-cell vesicles in this study (blue) with the previously reported TRPM4 structure obtained in LMNG detergent (PDB 9MTA; cyan) from Teixeira-Duarte *et al*. Side and top views are shown^13^. The two structures superimpose with a root mean square deviation (RMSD) of 1.895 Å. **(c)** Representative 2D class averages of TRPM4 particles derived from whole-cell vesicles prepared at different temperatures. Upper panel, vesicles prepared at 8 °C showing well-defined structural features and readily identifiable TRPM4 densities. Lower panel, vesicles prepared at 37 °C in the presence of CaCl₂ showing blurred and poorly resolved structural features, indicative of increased particle heterogeneity and preventing high-resolution structure determination.

**Supplementary Figure 3:**
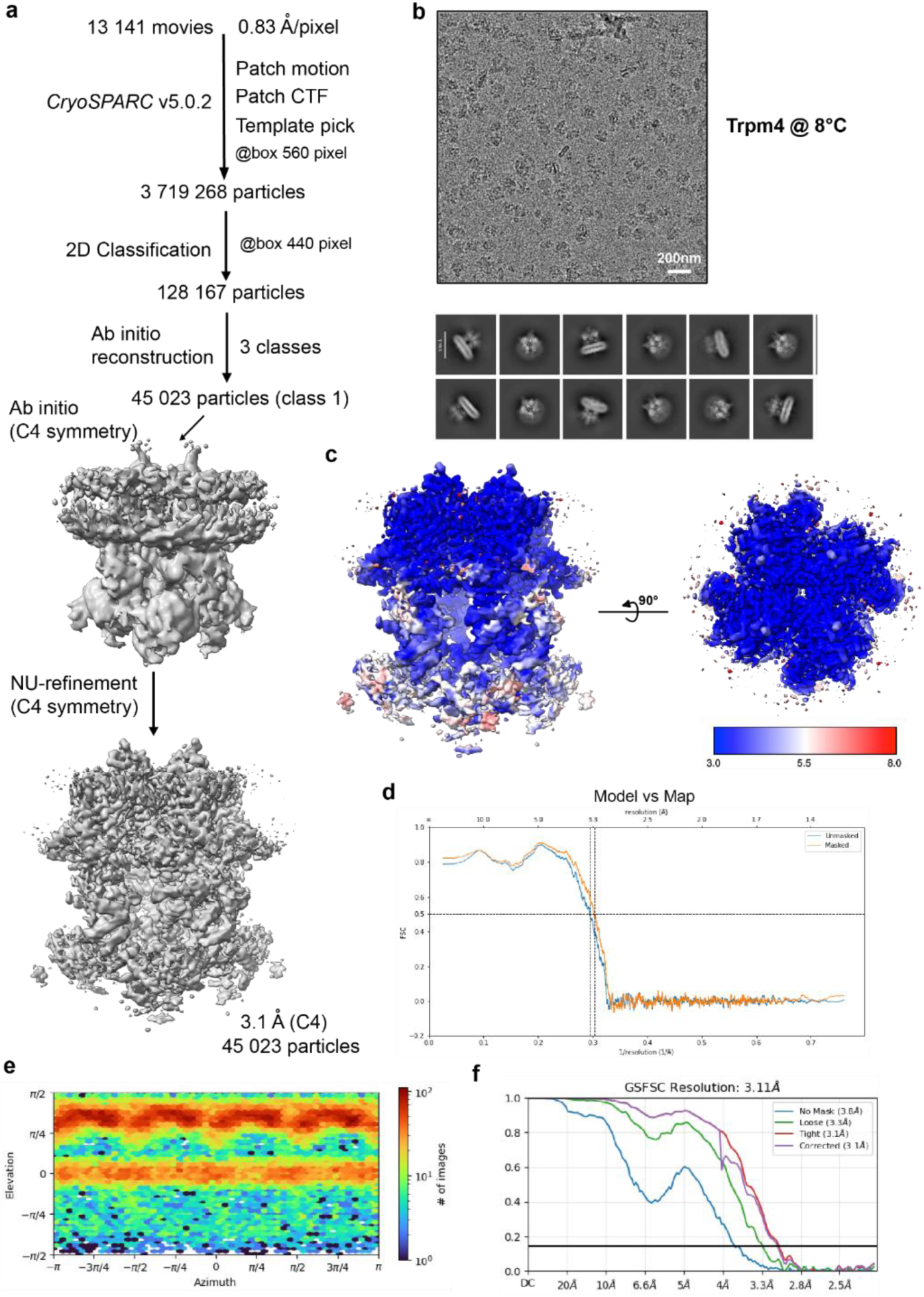
Cryo-EM data processing workflow and map resolution. (a) The image processing workflow of TRPM4_apo, 8 °C_. (b) micrograph and 2D classes (c) Local resolution (d) Model vs map FSC curves. (e) particle direction distribution. (f) FSC curves.

**Supplementary Figure 4:**
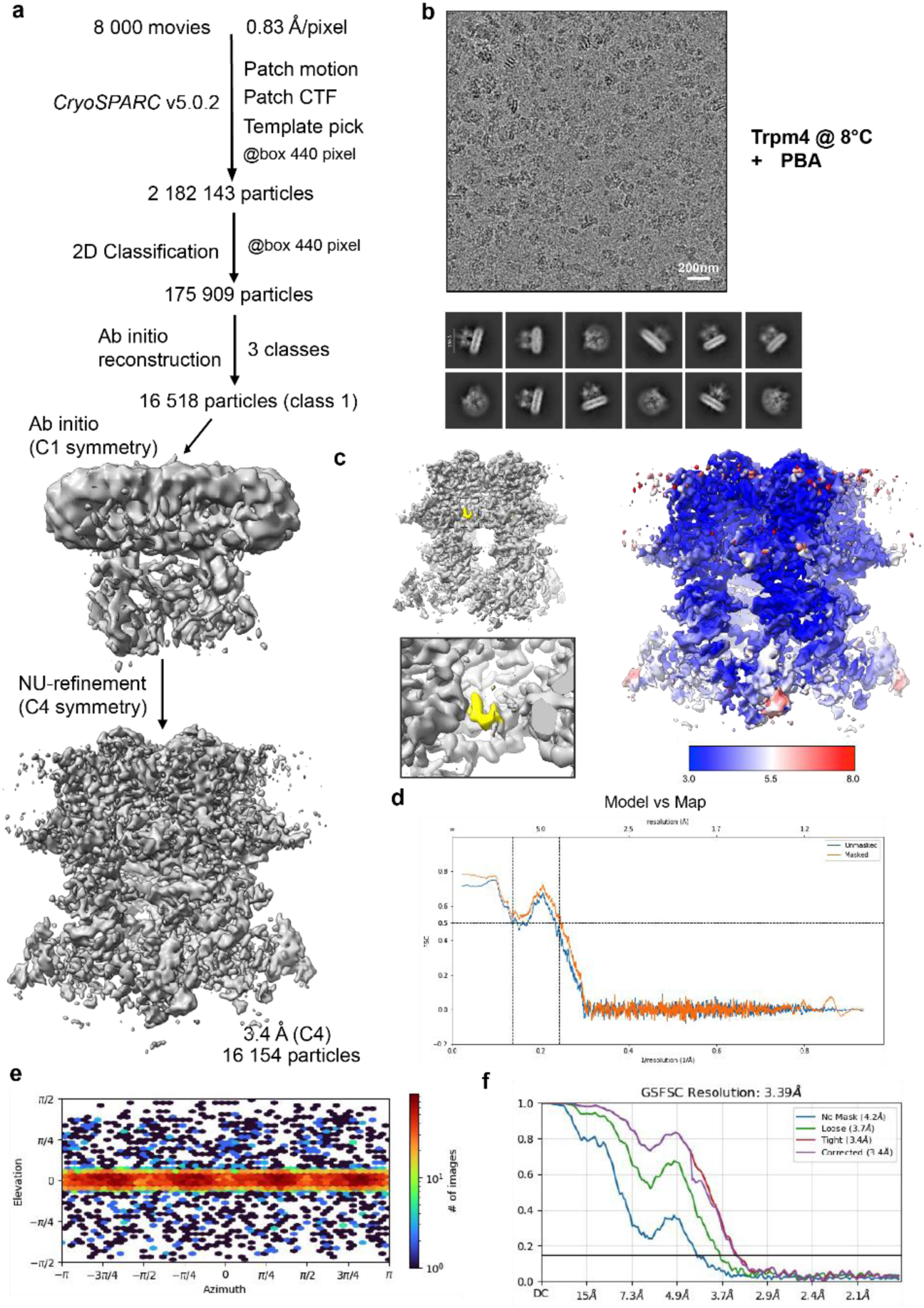
Cryo-EM data processing workflow and map resolution. (a) The image processing workflow of TRPM4_PBA, 8 °C_. (b) micrograph and 2D classes (c) Local resolution and PBA density (yellow) (d) Model vs map FSC curves. (e) particle direction distribution. (f) FSC curves.

**Supplementary Figure 5:**
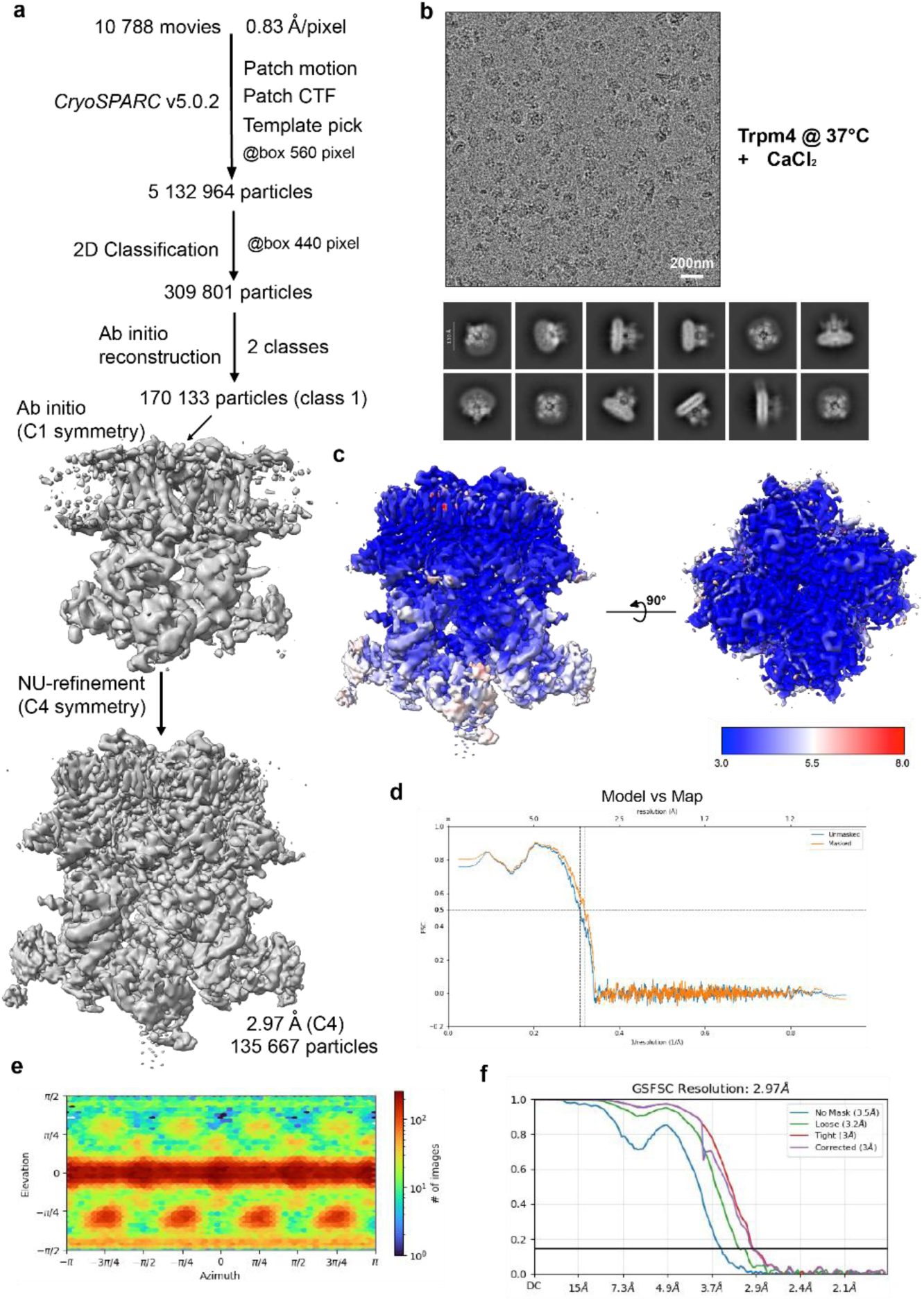
Cryo-EM data processing workflow and map resolution. (a) The image processing workflow of TRPM4_apo, 37 °C_. (b) micrograph and 2D classes (c) Local resolution (d) Model vs map FSC curves. (e) particle direction distribution. (f) FSC curves.

**Supplementary Figure 6:**
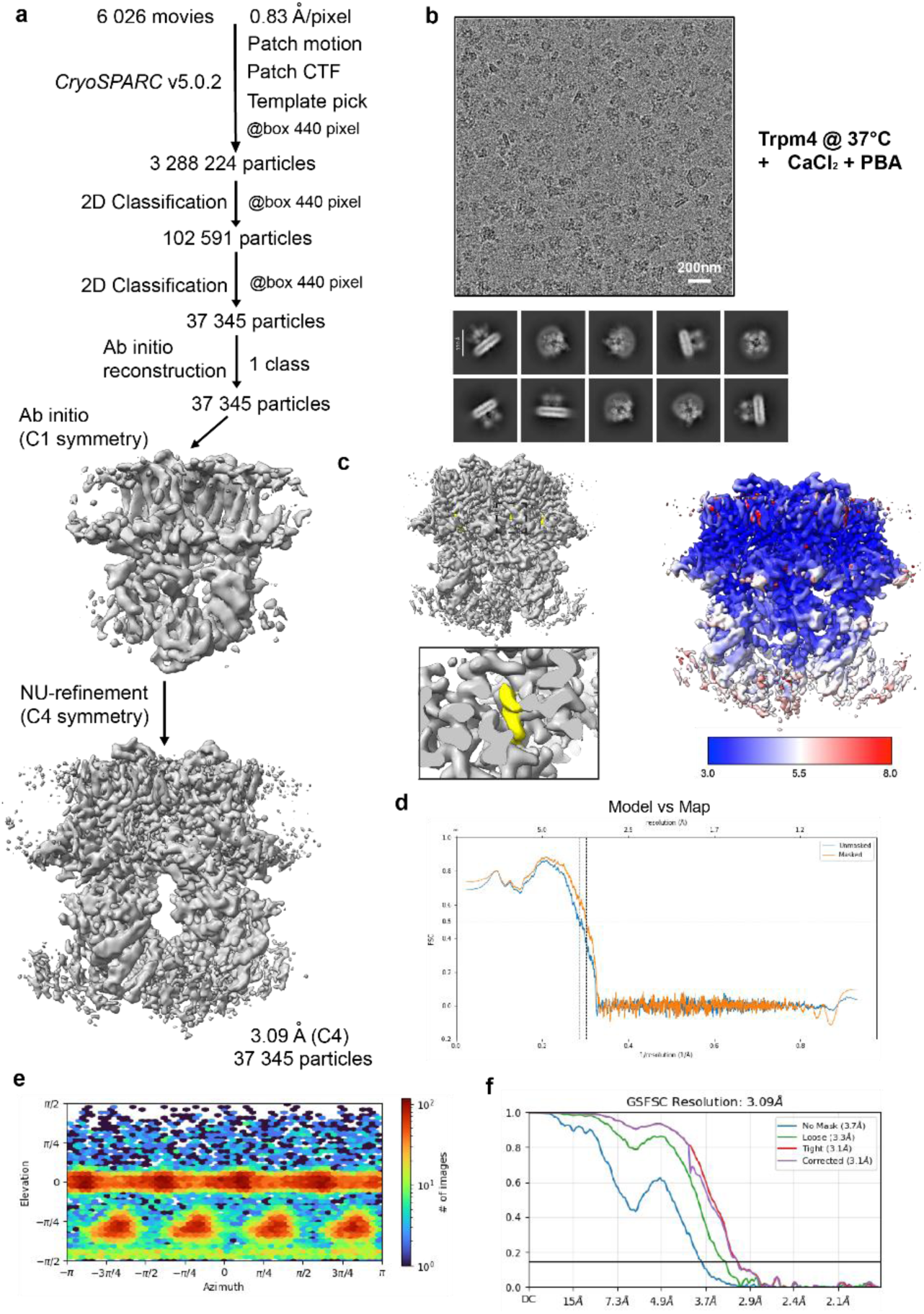
Cryo-EM data processing workflow and map resolution. (a) The image processing workflow of TRPM4_PBA, 37 °C_. (b) micrograph and 2D classes (c) Local resolution and PBA density (yellow) (d) Model vs map FSC curves. (e) particle direction distribution. (f) FSC curves.

**Supplementary Figure 7:**
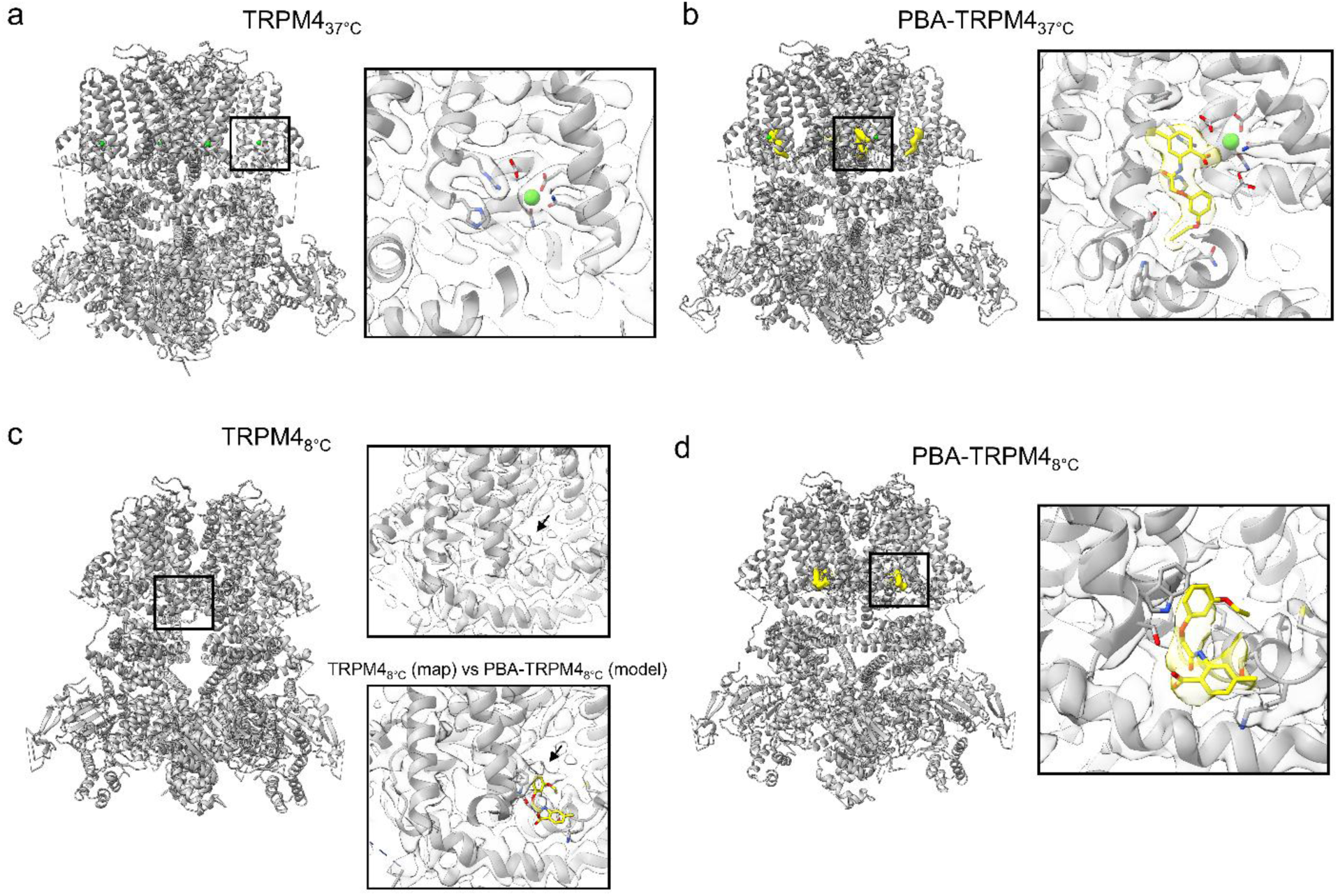
Cryo-EM density surrounding PBA-binding sites and neighbouring residues. **(a)** TRPM4 at 37 °C in the absence of PBA. The boxed region highlights the PBA-binding region observed in the ligand-bound structure. The enlarged view shows density for neighbouring residues and the bound Ca²⁺ ion, but no corresponding density for PBA. **(b)** PBA-bound TRPM4 at 37 °C. The enlarged view shows the cryo-EM density surrounding PBA and neighbouring residues within the S1–S4 binding site. **(c)** TRPM4 at 8 °C in the absence of PBA. The upper enlarged view shows a non-protein density close to the 8 °C PBA-binding pocket. The lower enlarged view shows an overlay with the PBA-bound model, indicating that this density does not overlap with PBA and likely corresponds to a lipid-like species. **(d)** PBA-bound TRPM4 at 8 °C. The enlarged view shows the cryo-EM density surrounding PBA and neighbouring residues within the transmembrane inhibitor-binding pocket.

